# Inducible and reversible phenotypes in a novel mouse model of Friedreich’s Ataxia

**DOI:** 10.1101/137265

**Authors:** Vijayendran Chandran, Kun Gao, Vivek Swarup, Revital Versano, Hongmei Dong, Maria C. Jordan, Daniel H. Geschwind

## Abstract

Friedreich’s ataxia (FRDA), the most common inherited ataxia, is caused by recessive mutations that reduce the levels of frataxin (FXN), a mitochondrial iron binding protein. We developed an inducible mouse model of *Fxn* deficiency that enabled us to control the onset and progression of disease phenotypes by the modulation of *Fxn* levels. Systemic knockdown of *Fxn* in adult mice led to multiple phenotypes paralleling those observed in human patients across multiple organ systems. By reversing knockdown after clinical features appear, we were able to determine to what extent observed phenotypes represent reversible cellular dysfunction. Remarkably, upon restoration of near wild-type FXN levels, we observed significant recovery of function, associated pathology and transcriptomic dysregulation even after substantial motor dysfunction and pathology were observed. This model will be of broad utility in therapeutic development and in refining our understanding of the relative contribution of reversible cellular dysfunction at different stages in disease.

## INTRODUCTION

A guanine-adenine-adenine (GAA) trinucleotide repeat expansion within the first intron of the frataxin (*Fxn*) gene in the chromosome 9 is the major cause of Friedreich’s ataxia (FRDA), the most commonly inherited ataxia(1, 2). Due to recessive inheritance of this GAA repeat expansion, patients have a marked deficiency of *Fxn* mRNA and protein levels caused by reduced *Fxn* gene transcription(2, 3). Indeed, the GAA expansion size has been shown to correlate with residual *Fxn* levels, earlier onset and increased severity of disease(4, 5). The prevalence of heterozygous carriers of the GAA expansion is between 1:60 and 1:110 in European populations, and the heterozygous carriers of a *Fxn* mutation are not clinically affected(6, 7).

FRDA is an early-onset neurodegenerative disease that progressively impairs motor function, leading to ataxic gait, cardiac abnormalities, and other medical co-morbidities, ultimately resulting in early mortality (median age of death, 35 years)(8, 9). However, identification of the causal gene led to identification of a significant number of patients with late onset, they tend to have slower progression with less severe phenotype and are associated with smaller GAA expansions(10). This shorter expansion enables residual *Fxn* expression(11), thus modifying the classical FDRA phenotype, consistent with other data indicating that *Fxn* deficiency is directly related to the FRDA phenotype(12). Extra-neurologic symptoms including metabolic dysfunction and insulin intolerance are observed in the majority and frank type I diabetes is observed in approximately 15% of patients, the severity of which is related to increasing repeat length(13-15). The mechanisms by which *Fxn* reduction leads to clinical symptoms and signs remain to be elucidated, but molecular and cellular dysfunction mediated by a critical reduction in *Fxn* levels plays a central role(16).

FXN is a nuclear-encoded mitochondrial protein involved in the biogenesis of iron-sulphur clusters, which are important for the function of the mitochondrial respiratory chain activity(17, 18). Studies in mouse have shown that *Fxn* plays an important role during embryonic development, as homozygous frataxin knockout mice display embryonic lethality(19), consistent with FXN’s evolutionary conservation from yeast to human(2, 3). Over the past several years, multiple mouse models of frataxin deficiency, including a knock-in knockout model(20), repeat expansion knock-in model(20), transgenic mice containing the entire *Fxn* gene within a human yeast artificial chromosome, YG8R and YG22R(21, 22), as well as a conditional *Fxn* knockout mouse, including the cardiac-specific(23) and a neuron specific model(23) have been generated. These existing transgenic and heterozygous knockout FRDA animal models are either mildly symptomatic, or restricted in their ability to recapitulate the spatial and temporal aspects of systemic FRDA pathology when they are engineered as tissue-specific conditional knockouts(20-25). Despite advances made towards elucidating FRDA pathogenesis, many questions remain due to the need for mouse models better recapitulating key disease features to understand frataxin protein function, disease pathogenesis and to test therapeutic agents.

In this regard, one crucial question facing therapeutic development and clinical trials in FRDA is the reversibility of symptoms. The natural history of the disorder has been well described(26, 27), it is not known how clinical features such as significant motor disability relate to reversible processes (e.g. underlying neuronal dysfunction) or reflect irreversible cell death. It is often assumed that clinically significant ataxia and motor dysfunction reflects neurodegeneration, although this may not be the case. This issue, while critically important for therapeutic development, is difficult to address in patients, but we reasoned that we could begin to address this question in an appropriate mouse model.

Here we report an inducible mouse model for FRDA, the FRDAkd mouse, that permits reversible, and yet substantial frataxin knockdown, allowing detailed studies of the temporal progression, or recovery following restoration of frataxin expression - the latter permitting exploration of disease reversal given optimal treatment (normalization of *Fxn* levels). We observe that *Fxn* knockdown leads to behavioral, physiological, pathological and molecular deficits in FRDAkd mice paralleling those observed in patients, including severe ataxia, cardiac conduction defects and increased left ventricular wall thickness, iron deposition, mitochondrial abnormalities, low aconitase activity, and degeneration of dorsal root ganglia, retina, as well as early mortality. We identify a signature of molecular pathway dysfunction via genome-wide transcriptome analyses, and show reversal of this molecular phenotype and as well as behavioral and pathological measures, even in the setting of significant disability due to motor dysfunction in FRDAkd animals.

## RESULTS

### RNA interference based in vivo frataxin knockdown and rescue

To investigate the neurological and cardiac effects linked to reduced FXN levels and to create a model for testing new therapies *in vivo*, we sought to generate mice that develop titratable clinical and pathological features of FRDA. We employed recombinase-mediated cassette exchange for genomic integration of a single copy shRNA transgene (doxycycline-inducible) that can mediate frataxin silencing temporally under the control of the H1 promoter via its insertion in a defined genomic locus that is widely expressed (rosa26 genomic locus)(28). This allowed us to circumvent the lethal effect of organism-wide knockout, while permitting significant frataxin reduction in all tissues. Depending on the dose of the inductor doxycycline (dox), temporal *Fxn* knockdown was achieved to control the onset and progression of the disease (Fig. 1a).

**Figure 1:**
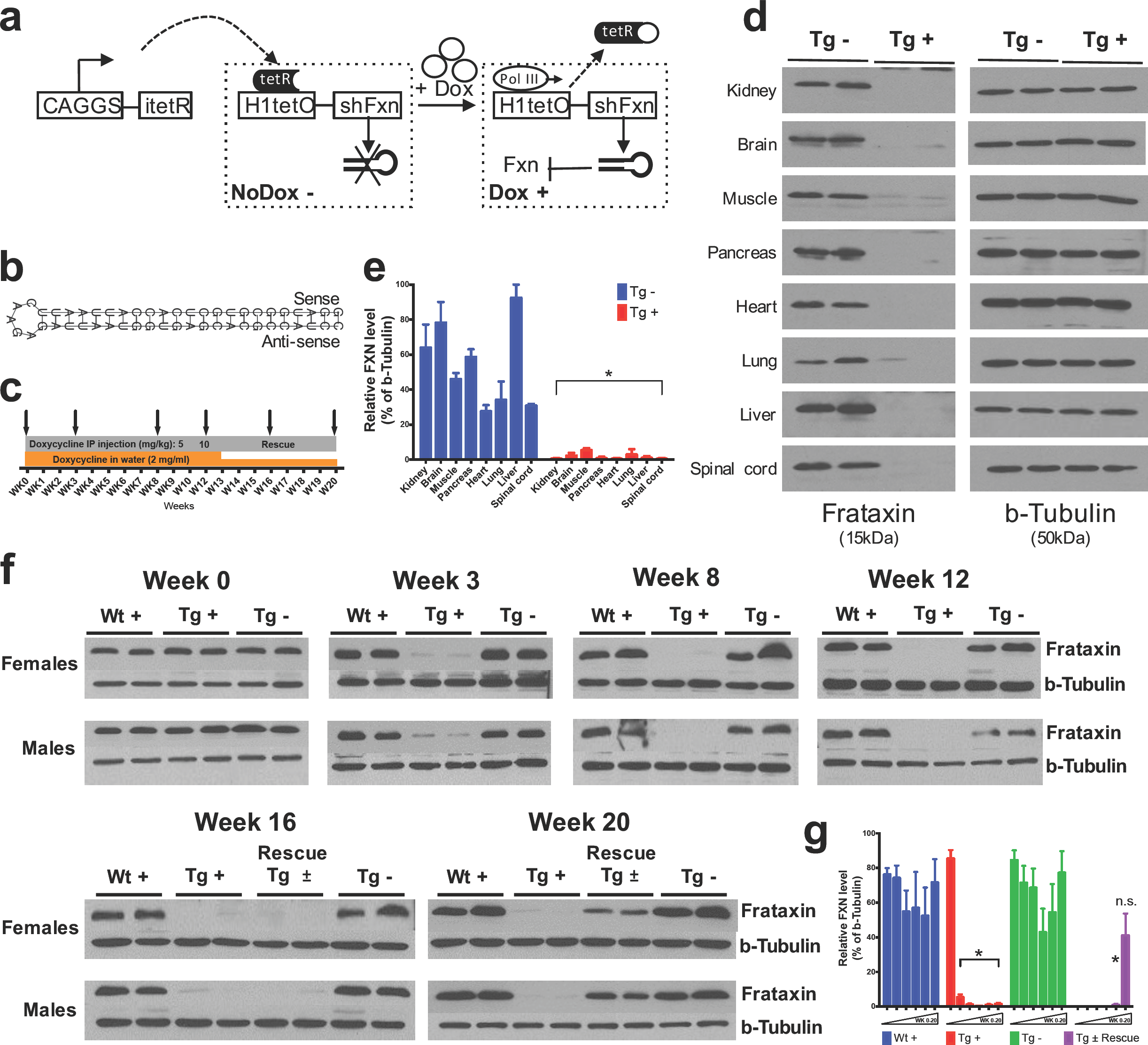
Efficient temporal in vivo frataxin knockdown and rescue. (**a**) Schematic representation of the inducible vector system to mediated expression of shRNA expression against frataxin. The vector contains shRNA sequence against frataxin (*Fxn*) gene regulated by the H1 promoter with tet-operator sequences (tetO) and tet repressor (tetR) under the control of the CAGGS promoter. Transcription of the *Fxn* shRNA is blocked in cells expressing the tetR. Upon induction by doxycycline (dox), tetR is removed from the tetO sequences inserted into the promoter, allowing transcription of shRNA against *Fxn*. shRNA expression leads to RNAi-mediated knockdown of *Fxn* gene. (**b**) Predicted minimum free energy secondary structure of expressed shRNA targeting the *Fxn* is shown with the sequence (sense strand) and its complement sequence (antisense strand) in the duplex form along with hairpin loop. (**c**) Timeline for doxycycline treatment of mice with a double IP injection (5 and 10 mg/kg) of dox per week and in drinking water (2 mg/ml). IP injections and dox in water was withdrawn for rescue animals. Arrow signs indicated different time intervals considered for downstream analyses. (**d**) Transgenic mice without (Tg -) or with (Tg +) doxycycline treatment (see c) for 20 weeks were analyzed by Western blot for protein levels of FXN in various organs. (**e**) For quantification, FXN values were normalized to the level of btubulin in each lane. (**f**) Time series FXN knockdown and rescue in liver. Wild-type mice with dox (Wt +), transgenic mice with (Tg +) and without (Tg -) dox, and transgenic mice with (Tg ±) dox removal (rescue) samples were analyzed for week 0,3,8,12,16 and 20. Rescue animals (Tg ±) were given dox for 12 weeks and doxycycline was withdrawn for additional 4 and 8 weeks. (**g**) For quantification, FXN values were normalized to the level of b-tubulin in each lane. N=4 animals per group, *=p<0.01; two-way ANOVA test; n.s., not significant. Error bars represent mean ± SEM for panels e and g.

Six different shRNA sequences were screened in vitro to obtain a highly efficient shRNA targeting the mature coding sequence of frataxin (Fig. 1b and **Supplementary Fig. 1**). Transgenic animals (FRDAkd) containing a single copy of this efficient shRNA transgene (Fig. 1b) were generated and characterized. First, to test *Fxn* knockdown efficiency, we explored the response to dox at varying escalating doses in drinking water. At the higher doses, we observed mortality as early as two weeks and a 100% mortality rate by 5 to 6 weeks, not permitting extended time series analyses (Online Methods). We found that the combination of 2 mg/ml in drinking water coupled with 5 or 10 mg/kg intraperitoneal injection of dox twice per week, led to efficient *Fxn* knockdown within two weeks post treatment initiation, while avoiding a high early mortality rate (Online Methods). Hence, for all subsequent experiments we utilized this regimen to model the chronicity of this disorder in patients by balancing the gradual appearance of clinical signs and decline in function, while limiting early demise (Fig. 1c).

To determine the effect of *Fxn* deficiency in adult mice after a period of normal development (similar to a later onset phenotype in humans), which would allow establishment of stable baselines, and to obtain relatively homogeneous data from behavioral tests(29) we initiated dox at 3 months. Following twenty weeks with (Tg +) or without (Tg -) doxycycline administration (Fig. 1c; Online Methods), we observed highly efficient silencing of *Fxn*, reaching greater than 90% knockdown across multiple CNS and non-CNS tissues (P < 0.05, two-way ANOVA; Fig. 1d, e). Using this regimen, time series western blot analyses of 80 independent animals (wildtype with dox (Wt +) N= 24; transgenic with dox (Tg +) N= 24; transgenic without dox (Tg -) N= 24; transgenic with dox removal (Rescue Tg ±) N= 8, at weeks 0, 3, 8, 12, 16 and 20) confirmed efficient silencing as early as 3 weeks, and efficient rescue, as evidenced by normal frataxin levels, post 8 weeks dox removal (P < 0.05, two-way ANOVA; Fig. 1f, g). Together, the results indicate that FRDAkd mice treated with dox are effectively FXN depleted in a temporal fashion and that *Fxn* expression can be reversed efficiently by dox removal, making it suitable for studying pathological and clinical phenotypes associated with FRDA.

### Frataxin knockdown mice exhibit neurological deficits

The major neurologic symptom in FRDA is ataxia, which in conjunction with other neurological deficits including axonal neuropathy and dorsal root ganglion loss, contributes to the gait disorder and neurological disability(8, 9). So, we first determined whether *Fxn* knockdown impacted the behavior of Tg and Wt mice with (+) or without (-) dox, or with dox, followed by its removal (±; the “rescue” condition), leading to a total of five groups subjected to a battery of motor behavioral tasks (each group N = 15-30; total mice = 108; Online Methods) (Fig. 2a-g). Wt - and Wt + control groups were included to access baseline and to have a control for any potential dox effect on behavior; Tg - and Tg + transgenic groups were compared to examine the effect of genotype and *Fxn* knockdown on behavior; The Tg ± group was included to study the effect of FXN restoration after knockdown (rescue). We observed significant weight loss and reduced survival ratio (<90%) at 25 weeks with dox treatment in *Fxn* knockdown animals (Tg +) when compared to other control groups (P < 0.05, two-way ANOVA; Fig. 2a,b). Tg + mice exhibited a shorter distance travelled at both 12 and 24 weeks in comparison to control animals, consistent with decreased locomotor activity (P < 0.05, two-way ANOVA; Fig. 2c). Next, we assessed gait ataxia(9) using paw print analysis(30) (Online Methods). The *Fxn* knockdown mice (Tg +) displayed reduced hind and front limb stride length when compared with Tg-, as well as the Wt control + and - dox at 12 and 24 weeks, suggesting ataxic gait (P < 0.05, two-way ANOVA; Fig. 2d,e and **Supplementary Fig. 2**). Grip strength testing also showed that Tg + animals displayed defect in their forelimb muscular strength at 12 and 24 weeks when compared with other groups (P < 0.05, two-way ANOVA; Fig. 2f). Finally, motor coordination and balance were evaluated using the Rotarod test. Whereas no significant difference in time spent on the rod before falling off was seen between Wt + or Wt - or Tg - and Tg ± mice after 12 weeks post dox removal (rescue), chronically treated mice (Tg + mice) from 12 weeks onward fell significantly faster, indicative of motor impairments (P < 0.05, two-way ANOVA; Fig. 2g). These observations suggest that the knockdown of *Fxn* in mice causes motor deficits indicative of ataxia similar to FRDA patients(9), and demonstrates the necessity of normal levels of *Fxn* expression in adults for proper neurological function.

**Figure 2:**
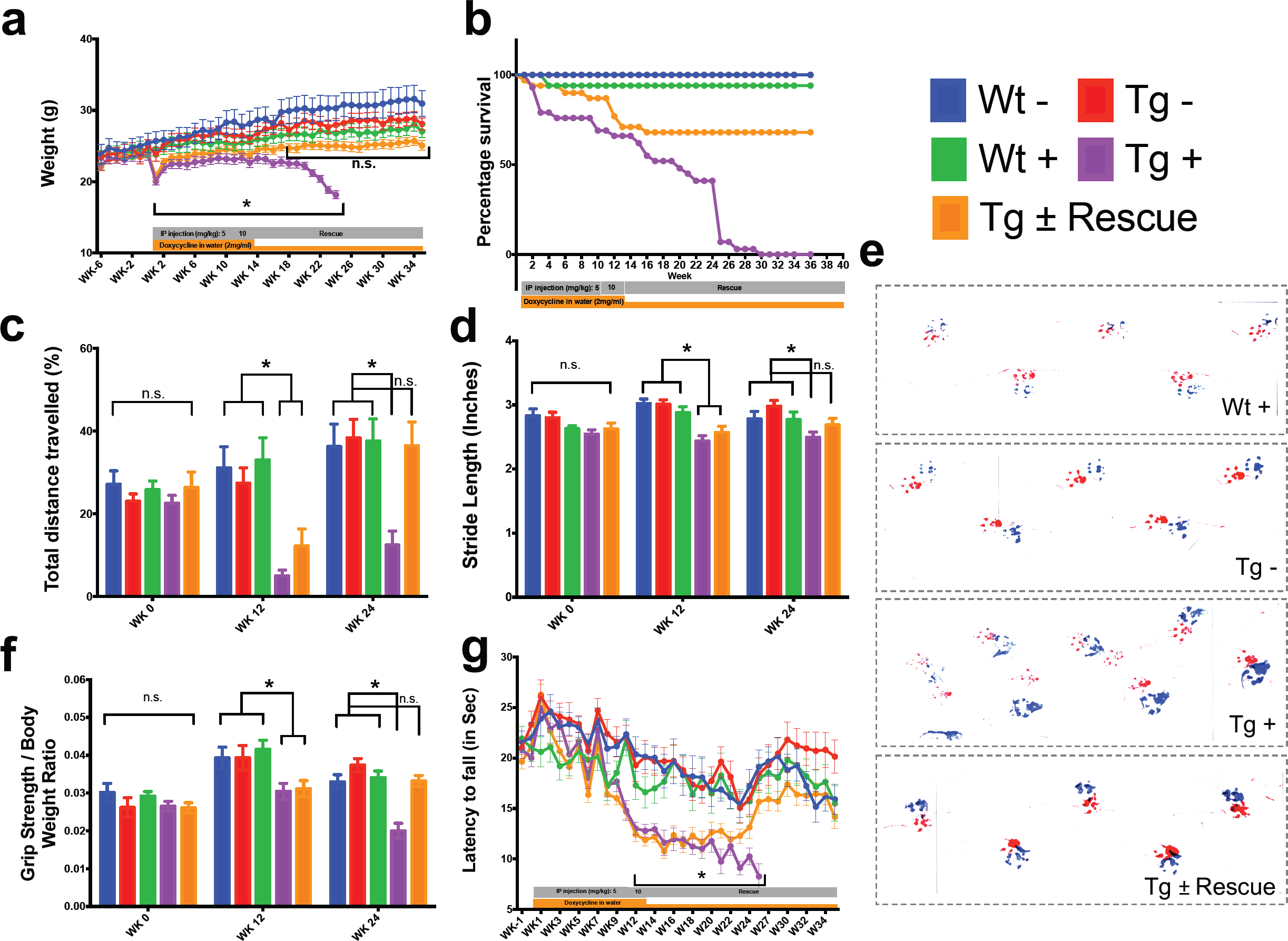
Neurological deficits due to frataxin knockdown. Body weight, survival, open field, gait analysis, grip strength and rotarod in five groups of animals; wild-type mice with dox (Wt +, n= 16(WK 0), n= 16(WK 12), n= 16(WK 24)) and without dox (Wt -, n= 16(WK 0), n= 16(WK 12), n= 16(WK 24)), transgenic mice with dox (Tg +, n= 30(WK 0), n= 21(WK 12), n= 15(WK 24)) and without dox (Tg -, n= 16(WK 0), n= 15(WK 12), n= 15(WK 24)), and transgenic mice with dox removal (Tg ± Rescue, n= 30(WK 0), n= 21(WK 12), n= 20(WK 24)). (**a**) Body weight from before 6 weeks and upto 34 weeks after dox treatment. (**b**) Survival was significantly declined in dox treated Tg + group of animals, no mortality was observed after dox withdrawal (Tg ± Rescue). (**c**) Open field test showing significant decline in total distance traveled by the dox treatment transgenic animals (Tg +) at 12 and 24 weeks when compared across all other groups. After dox withdrawal, there were no differences between the rescue group (Tg ± Rescue) and the three control groups at week 24. (**d**) Gait footprint analysis of all five groups of mice at 0, 12 and 24 weeks was evaluated for stride length. Dox treated transgenic (Tg +) animals revealed abnormalities in walking patterns displaying significantly reduced stride length, however the rescue group (Tg ± Rescue) displayed normal stride length when compared to other groups. (**e**) Representative walking footprint patterns. (**f**) Grip-strength test, dox treated transgenic (Tg +) mice had reduced forelimb grip strength at 12 and 24 weeks when compared across all other groups. After dox withdrawal, there were no significant differences between the rescue group (Tg ± Rescue) and the three control groups at week 24. (**g**) Rotarod test in mice upto 34 weeks after dox treatment. Dox treated transgenic (Tg +) animals stayed less time on the rotarod than the control groups, although after dox withdrawal, there was no significant difference between the rescue group (Tg ± Rescue) and the three control groups. Values between all five groups are shown as mean ± SME. Two-way ANOVA test *=P≤0.001; n.s., not significant.

### Frataxin knockdown leads to cardiomyopathy

Cardiac dysfunction is the most common cause of mortality in FRDA(31, 32). To examine impaired cardiac function in FRDAkd knockdown animals, we employed electrocardiogram (ECG) and echocardiogram analyses to measure electrical activity and monitor cardiac dimensions. The ECG of Tg + mice displayed a significant increase in QT interval duration when compared to the other control/comparison groups at both 12 and 24 weeks post dox treatment, suggesting abnormal heart rate and rhythm (arrhythmia)(33) (Online Methods; P < 0.05, two-way ANOVA; Fig. 3a,e). However, rescue animals (Tg ±) 12 weeks after dox removal showed normal electrocardiogram, demonstrating that by restoring the FXN levels, the prolonged QT interval can be reversed (P < 0.05, two-way ANOVA; Fig. 3b,e). We also observed that the Tg + animals at week 24 displayed absence of P-waves suggesting an atrial conduction abnormality(34) (Fig. 3c), not observed in the rescued animals. Similar abnormalities have been variably observed in many FRDA patients(35).

**Figure 3:**
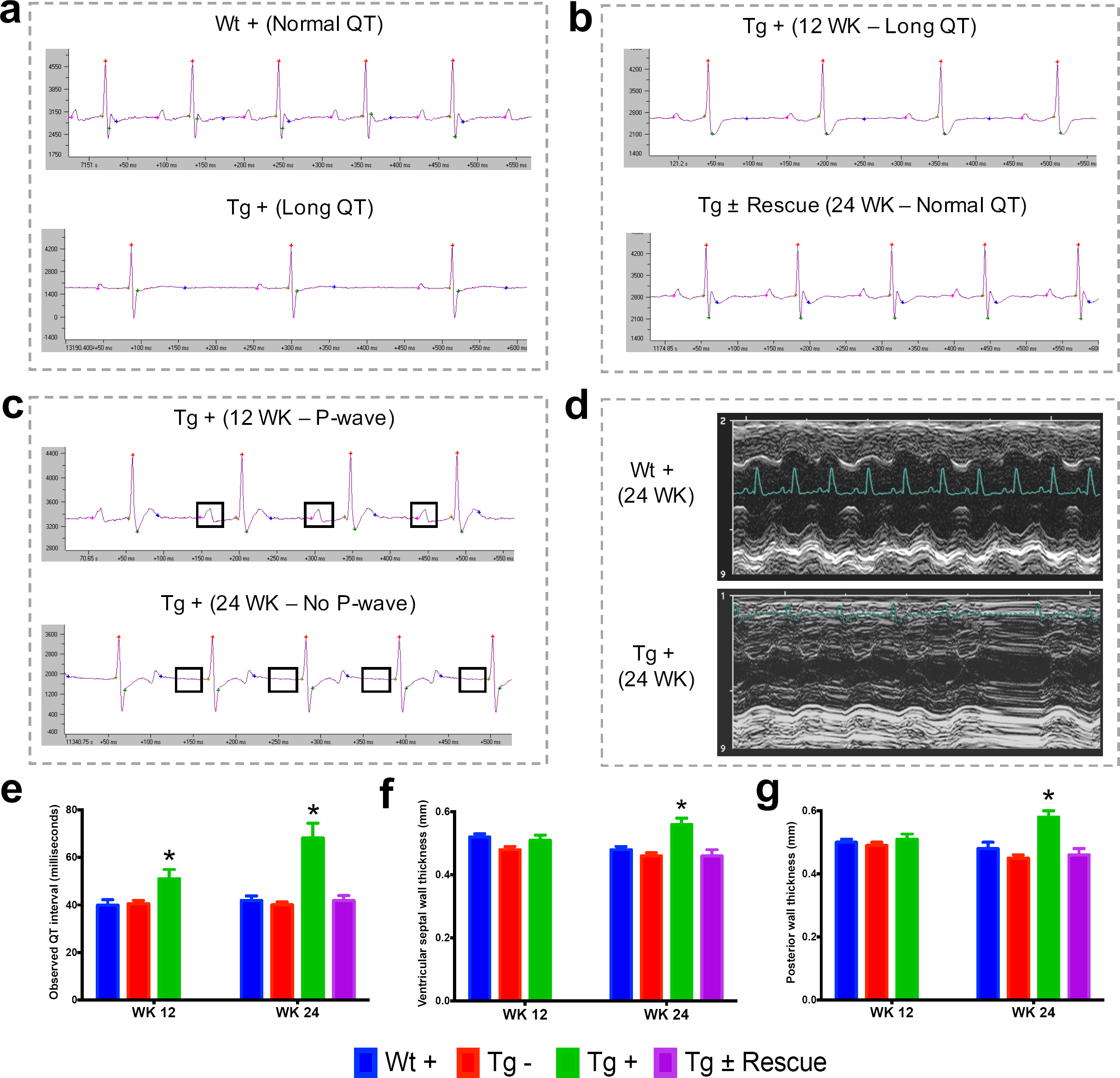
Frataxin knockdown mice exhibit signs of cardiomyopathy. (**a**) Representative traces of ECG recording in a wild-type and transgenic animal after dox treatment for 20 weeks, showing long QT intervals in Tg + animal. (**b**) ECG recording of a same dox treated transgenic (Tg +) animal at 12 and after dox withdrawal for additional 12 weeks (Tg ± Rescue), showing normal QT interval. (**c**) ECG recording of a same dox treated transgenic (Tg +) animal at 12 and 24 weeks, showing absences of P-wave only at 24 week. (**d**) Representative echocardiograms from the left ventricle of a wild-type and transgenic mouse at 24 week after dox treatment. (**e-f**) Quantification of observed QT interval (**e**), ventricular septal wall thickness (**f**) and posterior wall thickness (**g**) are shown for week 12 and 24. N = 6-8 animals per group, *=p<0.05; Student’s t test. Error bars represent mean ± SEM.

Progressive hypertrophic cardiomyopathy (thickening of ventricular walls) related to the severity of frataxin deficiency(35-37) is frequently observed in FRDA patients(31). To confirm structural cardiac abnormalities, echocardiogram analyses were performed, focusing on left ventricular function. At 12 weeks there was a non-significant trend towards increasing ventricular wall thickness. However, by 24 weeks, dox treated transgenic animals (Tg +) exhibited ventricular and posterior wall thickening, suggesting hypertrophic cardiomyopathy in these animals when compared to other control groups (P < 0.05, two-way ANOVA; Fig. 3d,f,g). Together, these observations indicate that the transgenic mice exhibit progressive cardiomyopathy due to reduced level of *Fxn*, supporting the utilization of Tg + mice for examining the molecular mechanisms downstream of *Fxn* deficiency responsible for cardiac defects.

### Cardiac pathology observed with frataxin knockdown

We next explored pathological consequences of FRDA knockdown in FRDAkd mice. In FRDA patients, reduced frataxin induces severe myocardial remodeling, including cardiomyocyte iron accumulation, myocardial fibrosis and myofiber disarray(9). Indeed, we observed substantially increased myocardial iron in Tg + mice, as evidenced by increased ferric iron staining (Fig. 4a and **Supplementary Fig. 3**) and the increased expression of iron metabolic proteins, ferritin and ferroportin at 20 weeks (Fig. 4b and **Supplementary Fig. 3**). Cardiac fibrosis is commonly found in association with cardiac hypertrophy and failure(38). Histological analysis by Masson’s trichrome staining revealed excessive collagen deposition in Tg + mice hearts at 20 weeks when compared to other control groups, suggesting cardiac fibrosis (Fig. 4c). Further examination of cardiomyocyte ultrastructure by electron microscopy in control mouse (Wt +) heart demonstrates normally shaped mitochondria tightly packed between rows of sarcomeres (Fig. 4d). In contrast, Tg + mice demonstrate severe disorganization, displaying disordered and irregular sarcomeres with enlarged mitochondria at 20 weeks (Fig. 4d). In a minority of cases, but never in controls, we observed mitochondria with disorganized cristae and vacuoles in Tg + mouse heart at 20 weeks, suggesting mitochondrial degeneration (Fig. 4e). Next, by examining aconitase which is a Fe-S containing enzyme whose activity is reduced in FRDA patients(39, 40), activities in Tg + and other control groups, we observed decreased aconitase activity in the Tg + mouse heart at 20 weeks. Together these observations suggest that the knockdown of *Fxn* in mice causes cardiac pathology similar to that observed in patients(32).

**Figure 4:**
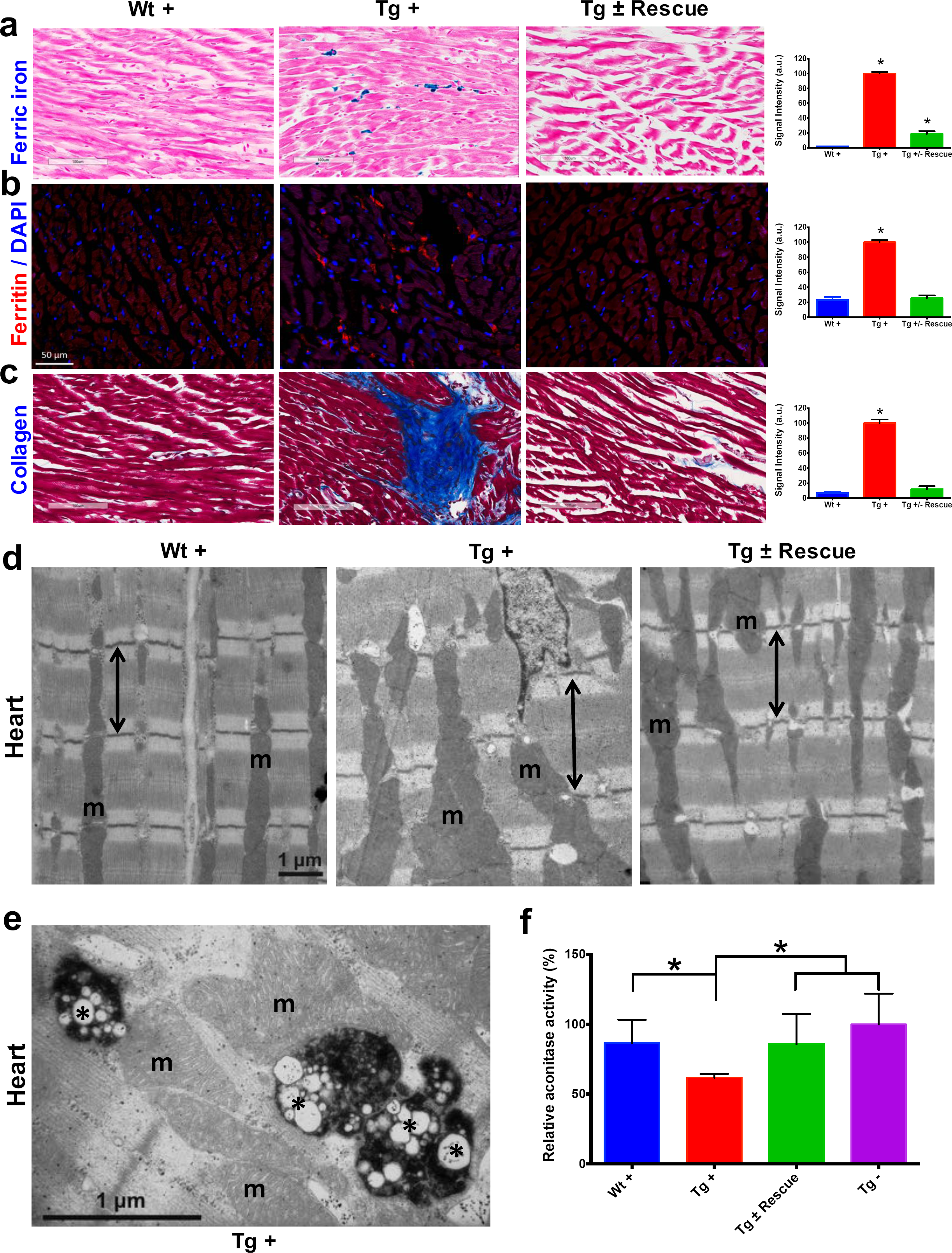
Cardiopathology of frataxin knockdown mice. (**a**) Gomori’s iron staining and quantification of iron deposition in dox treated transgenic (Tg +), wild-type (Wt +) and dox withdrawn transgenic (Tg ± Rescue) animals. Dox treated transgenic (Tg +) mice showing myocardial iron-overload (a) also displayed altered expression of ferritin protein (**b**) which is involved in iron storage. Both iron-overload and ferritin protein levels were significantly lower in Tg ± Rescue animals (a-b). (**c**) Masson’s trichrome staining and quantification showing increased fibrosis in Tg + mice when compared to Wt + and Tg ± Rescue animals. (**d**) Electron micrographs of cardiac muscle from Wt +, Tg + and Tg ± Rescue animals at 20 week after dox treatment. Double arrow lines indicate sarcomere. m = mitochondria. Scale bars, 1 μm. Data are representative of three biological replicates per group. (**e**) Higher magnification of electron micrographs of cardiac muscle from Tg + mice, showing normal (m) and degenerating mitochondria (asterisks). (**f**) Aconitase activity was assayed in triplicate in tissues removed from 3 hearts in each group. Values represent mean ± SME. One-way ANOVA test *=P≤0.05.

### Frataxin knockdown causes neuronal degeneration

In FRDA patients and mice with *Fxn* conditional knockout, a cell population that is seriously affected by frataxin reduction is the large sensory neurons of dorsal root ganglia (DRG), which results in their degeneration(9). As in heart tissue, lack of *Fxn* also induces mitochondrial dysfunction in the DRG(40). Thus, we assessed whether mitochondrial abnormalities are observed in Tg + mice DRG neurons by electron microscopy (Online Methods). As anticipated, no mitochondrial abnormalities were detected in DRG neurons in control groups, but a significant increase (P < 0.05, two-way ANOVA) in condensed mitochondria were detected in DRG neurons of Tg + mice at 20 weeks (Fig. 5a-c,e), but not in Tg - or Wt + mice. In most cases we observed lipid-like empty vacuoles associated with condensed mitochondria in DRG neurons of Tg + animals, but not in other control groups, suggesting neuronal degeneration, similar to what is observed in FRDA patients(9) (P < 0.05, two-way ANOVA; Online Methods; Fig. 5b,c). Previous results in human post-mortem spinal cords of FRDA patients showed a decrease in axon size and myelin fibers(8). By analyzing more than 2000 axons per group in the lumbar spinal cord cross-section of high-resolution electron micrographs, we observed significant reduction in axonal size and myelin sheath thickness in the spinal cord samples of Tg + mice when compared to other control groups at 20 weeks (P < 0.05, two-way ANOVA; Fig. 5f,g). In the cerebellum, the number of Purkinje cells, the cerebellar granular and molecular layers were not altered due to *Fxn* knockdown at 20 weeks (**Supplementary Fig. 4**).

**Figure 5:**
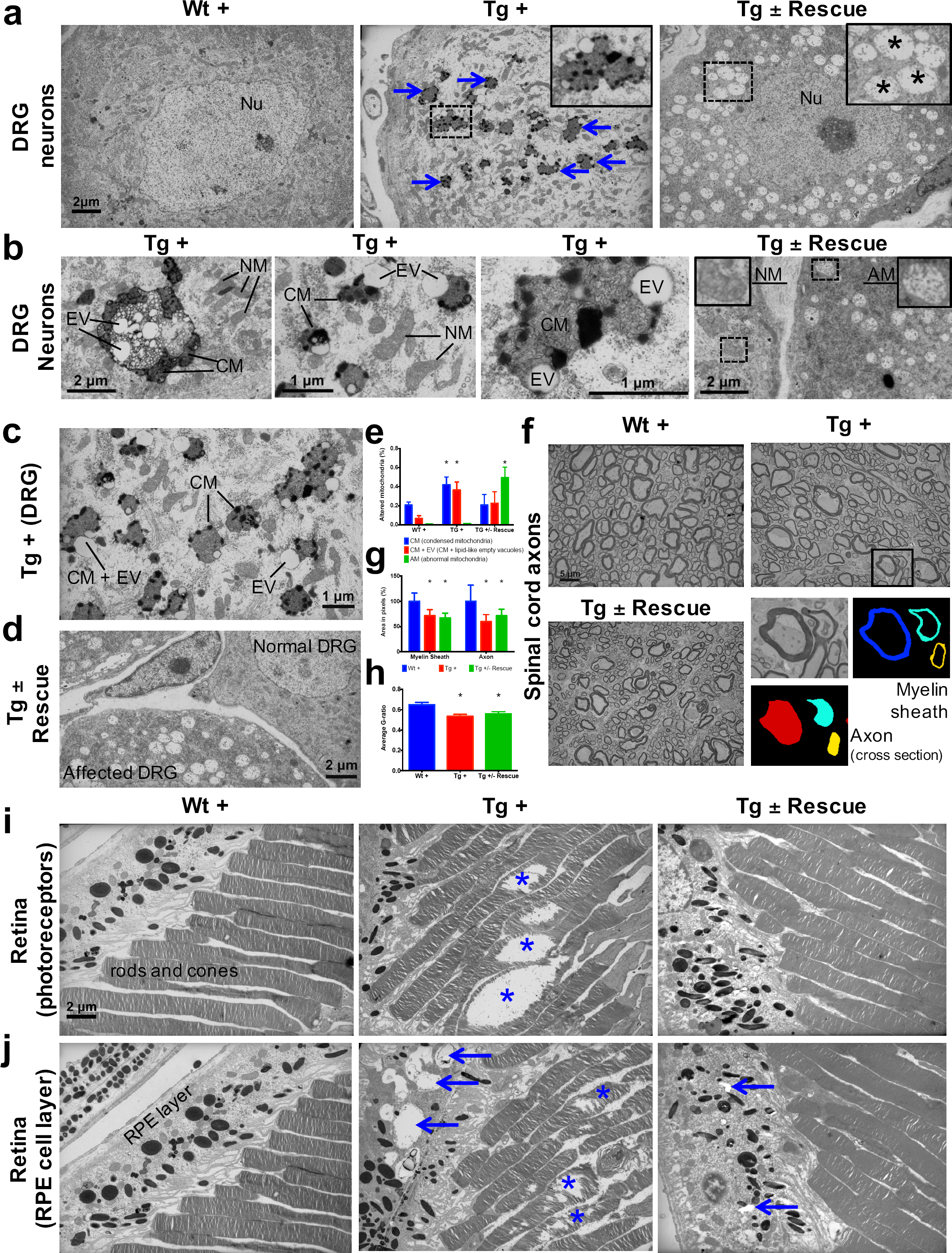
Frataxin knockdown mice exhibit neuronal degeneration. (**a**) Electron microscopic analysis of Wt +, Tg + and Tg ± Rescue animal DRG neurons at 20 week after dox treatment. Arrows indicate condensed mitochondria with lipid-like empty vacuoles. Insert in Tg + panel shows higher magnification of electron micrographs of condensed mitochondria with lipid-like empty vacuoles in DRG neurons. Insert in Tg ± Rescue panel shows higher magnification of abnormal mitochondria (asterisks). Nu= nucleus. (**b**) Higher magnification of electron micrographs of Tg + and Tg ± Rescue animal DRG neurons at 20 week after dox treatment. Tg + panel shows degenerating mitochondria and condensed mitochondria with lipid-like empty vacuoles in DRG neurons. Tg ± Rescue panel shows two DRG neurons, one consisting of normal mitochondria (insert) and the other neuron with abnormal mitochondria (insert). (**c**) Higher magnification of Tg + animals showing condensed mitochondria with lipid-like empty vacuoles in DRG neurons. (**d**) Higher magnification of Tg ± rescue animals showing two DRG neurons, one consisting of normal mitochondria (normal DRG) and the other neuron with abnormal mitochondria (affected DRG). NM= normal mitochondria, CM= condensed mitochondria, EV= lipid-like empty vacuoles, AM= abnormal mitochondria. (**e**) Quantification of altered mitochondria in DRG neurons. Data are from multiple images from three biological replicates per group. Values represent mean ± SME. Two-way ANOVA test *=P≤0.05. (**f**) Electron micrographs of spinal cord axon cross-section, displaying reduced myelin sheath thickness and axonal cross-section area in Tg + and Tg ± Rescue animals. Bottom panel shows representative area utilized for quantification. (**g-h**) Quantification of myelin sheath thickness and axonal cross-section area in the spinal cord. Data are from 2000 or more axons per group in the lumbar spinal cord cross-section of high-resolution electron micrographs from three biological replicates per group. Values represent mean ± SME. One-way ANOVA test *=P≤0.05. (**i**) Electron micrographs of rod and cone photoreceptor cells, showing their disruption in Tg + animals (asterisks). (**j**) Retinal pigment epithelium cell layer showing the presences of large vacuoles (arrows) in Tg + animals along with disruption in their photoreceptor cells (asterisks).

Next we explored the visual system, because in FRDA patients, various visual field defects have been reported, suggesting that photoreceptors and the retinal pigment epithelium (RPE) can be affected(41, 42). By electron microscopic examination of the photoreceptors in the retina, which are specialized neurons capable of phototransduction, we observed disruption in Tg + mice at 20 weeks (Online Methods; Fig. 5h). Previous work has shown that the disruption of photoreceptors is related to visual impairment(43). Similarly, we also found a significant increase in degenerating RPE cells with vacuoles in Tg + mice, which is involved in light absorption and maintenance of visual function (Fig. 5i). Together, these results suggest that *Fxn* knockdown in the CNS of adult mice results in degeneration of retinal neurons.

### Gene expression changes due to frataxin knockdown

Given the phenotypic parallels in the cardiac and nervous system abnormalities in FRDAkd mice with chronic *Fxn* reduction following treatment with dox, we next sought to explore genome wide molecular mechanisms and determine which pathways were affected in the heart and nervous system, and if they were reversible. We analyzed global gene expression profiles in the heart, cerebellum and DRGs from Tg +, Tg -, Wt + and Tg ± mice from 0, 3, 12, 16, 20 and plus 4, 8 weeks post dox treatment (n= 192). Differential expression analyses (Online Methods) identified 1959 genes differentially expressed in Tg + mice heart when compared to Wt + and Tg - mice (FDR < 5%, **Fig. 6a, Supplementary Table 1**). Similarly, we observed 709 and 206 genes differentially expressed in cerebellum and DRGs of Tg + mice. While cross tissue overlap in expression changes was significant, the majority of changes were tissue specific; only 31% and 38% of genes that were differentially expressed in the Tg + mice cerebellum and DRGs were also dysregulated in heart. Likewise, we observed a 19% overlap between DRGs and cerebellum, consistent with previous observations that *Fxn* reduction causes distinct molecular changes in different tissues(15).

**Figure 6:**
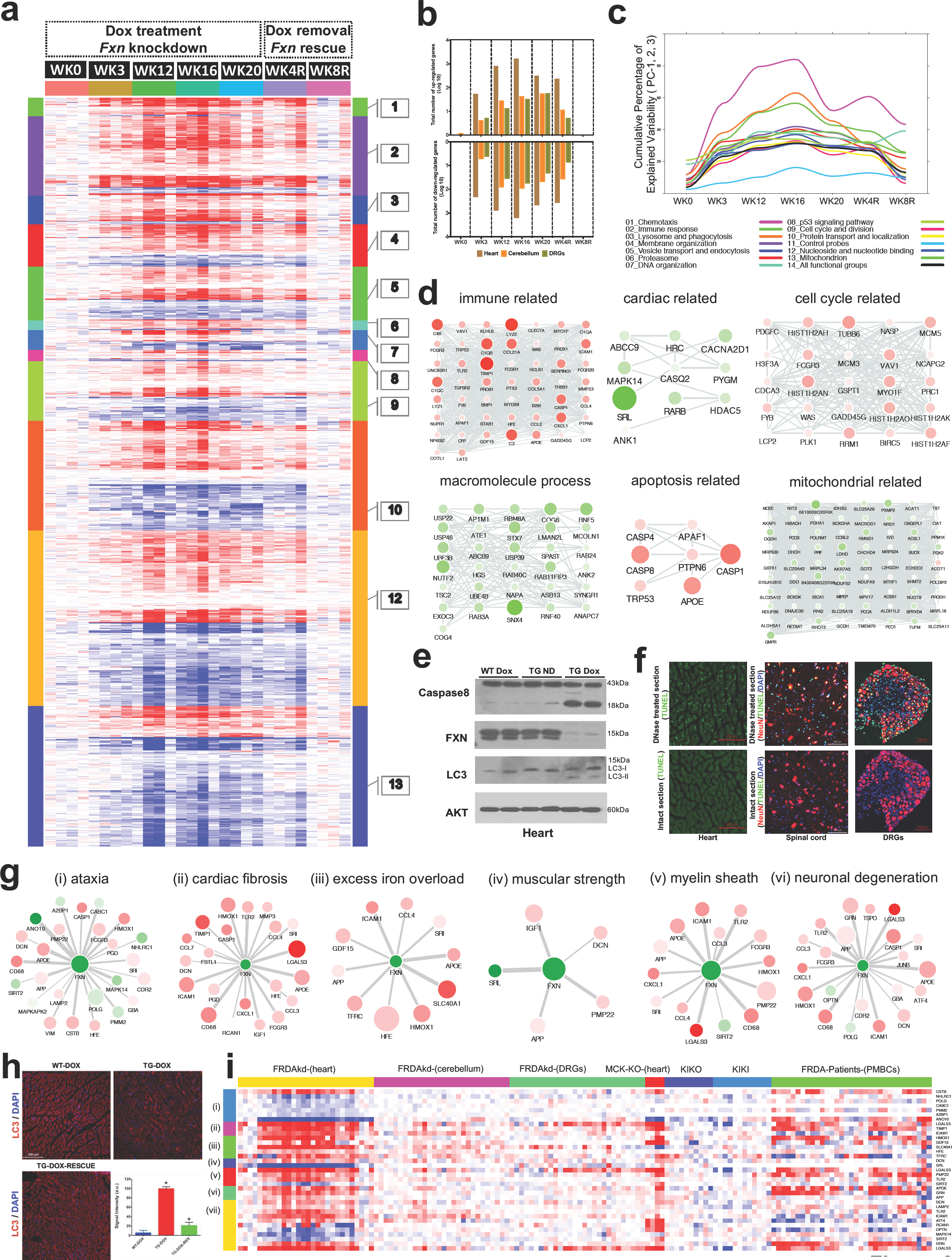
Gene expression analysis of frataxin knockdown mice. (**a**) Heat map of significantly up- and down-regulated genes (rows) in heart tissue of Tg + mice from 0, 3, 12, 16, 20 and plus 4, 8 weeks post dox treatment relative to controls are grouped into 13 functional categories. (**b**) Summary of differentially expressed genes during *Fxn* knockdown and rescue in heart, cerebellum and DRG tissues from four biological replicates. (**c**) Cumulative percent of variability in Tg + gene expression data explained by the first three principal component for each functional category. (**d**) Networks highlighting differentially expressed genes due to *Fxn* knockdown in Tg + mice for selected functional categories. Nodes represents genes and edges are present between nodes when their gene expression correlation is greater than 0.5. Node size and color (red= up-regulation and green= down-regulation) denotes extent of differential expression. (**e**) Western blot analyses of Caspase 8, FXN and LC3 following 20 weeks with or without dox treatment in heart. (**f**) TUNEL assay of heart, spinal cord and DRG neurons at 20 weeks after dox treatment in Tg + mice. (**g**) Literature associated gene networks highlighting differentially expressed genes due to *Fxn* knockdown in Tg + mice for selected key-terms. Nodes represents genes which has pairwise correlation greater than 0.5 with *Fxn* gene and edge size represents strength of their gene expression correlation. Node size correspond to number of PubMed hits with co-occurrence of gene and its corresponding key-term. Node color represents up-regulated (red) and down-regulated (green). (**h**) Representative images and quantification of LC3 staining in heart tissue at 20 weeks after dox treatment. Values represent mean from three biological replicates per group ± SME. One-way ANOVA test *=P≤0.05. (**i**) Heat map depicting expression of key genes (rows) across samples (columns) for seven groups (red corresponds to gene up-regulation and blue to down-regulation). Seven groups represents, (i) ataxia, (ii) cardiac fibrosis, (iii) excess iron overload, (iv) muscular strength, (v) myelin sheath, (vi) neuronal degeneration and (vii) autophagy related genes. Column represents, seven independent datasets obtained from, FRDAsh mice, cardiac specific knockout mice(23), knock-in knockout mice(20), knock-in mice(20) and patient peripheral blood mononuclear cells(44).

Next, we analyzed these differently expressed transcripts in cardiac tissue from Tg + mice with respect to cellular pathways. The top GO categories and KEGG pathways include chemotaxis, immune response, lysosome and phagocytosis, vesicle transport and endocytosis, p53 signaling pathway, cell cycle and division, protein transport and localization, nucleoside and nucleotide binding, and mitochondrion (Benjamini corrected P-value < 0.05) (**Fig. 6a, Supplementary Table 2**). To characterize the temporal patterns of these signaling cascades after frataxin knockdown and rescue, we examined their time course by PCA analyses of the gene expression profiles (Fig. 6b). By examining the cumulative explained variability of the first three principal components for these clusters of genes, we show that each of these functional groups are activated as early as three weeks after dox initiation (and remain elevated for up to 20 weeks); importantly, the aberrant expression of all of these clusters observed in Tg + mice are largely reversed after eight weeks of rescue via *Fxn* re-expression (Fig. 6a,b).

Another notable observation is that immune system activation is among the earliest pathways regulated after *Fxn* knockdown (Fig. 6b). This suggests that initiation of immune responses (innate and adaptive) is a direct consequence of *Fxn* knockdown. For example, 38 genes involved in the chemokine signaling pathway (KEGG: mmu04062) were significantly differentially expressed due to *Fxn* knockdown in Tg + mice heart (**Supplementary Fig. 5**). Cross validating these genes with previously published gene expression datasets obtained from FRDA patients(44) and mouse models associated with FRDA(20, 23), identified several genes involved in the chemokine signaling pathway (E.g.: CCL2, 3, 4, 7, CXCL1, 16, PRKCD, STAT3) consistently differentially expressed in these six independent FRDA associated datasets (**Supplementary Fig. 6**). This implicates a vital role for chemokines and immune response in FRDA pathology, as has been suggested for other neurodegenerative diseases(45-48).

### Shared and tissue specific gene expression changes

We next performed weighted gene co-expression network analysis (WGCNA)(49-51), a powerful method for understanding the modular network structure of the transcriptome, to organize the frataxin related transcriptional changes in an unbiased manner following knockdown and rescue. WGCNA permits identification of modules of highly co-expressed genes whose grouping reflects shared biological functions and key functional pathways, as well as key hub genes within the modules, and has been widely applied to understanding disease related transcriptomes(52, 53). By applying WGCNA and consensus network analysis(50) we identified 19 robust and reproducible co-expression modules (Online Methods; **Supplementary Fig. 7 and Supplementary Table 3**) between the three tissues datasets generated at 0, 3, 12, 16 and 20 weeks after *Fxn* knockdown and 4 and 8 weeks post dox removal. On the basis of the module eigengene correlation with time-dependent changes after *Fxn* knockdown, we first classified modules as up-regulated and down-regulated following knockdown (**Supplementary Fig. 8**). Next, based on the significant module trait relationships (Wilcoxon P-value < 0.05), we identified 11 modules strongly associated with *Fxn* knockdown: three down-regulated modules in two or more tissues after *Fxn* knockdown (yellow, lightgreen and turquoise) and three up-regulated modules (blue, purple, and black) (**Supplementary Fig. 8**), three down-regulated modules in heart, which were up-regulated in cerebellum (red, greenyellow and magenta) and two up-regulated modules in heart which were down-regulated in cerebellum (cyan and pink). Although six of the gene co-expression modules (yellow, lightgreen, turquoise, blue, purple, and black) in the heart, cerebellum and DRG following *Fxn* knockdown are highly preserved across tissues, five modules (red, greenyellow, magenta, cyan and pink) exhibit differential expression profiles suggesting tissue specific molecular changes, consistent with previous observations of shared and organ specific changes(15) (**Supplementary Fig. 8**).

As a first step toward functional annotation of the cross tissue modules, we applied GO and KEGG pathway enrichment analyses, which showed enrichment (Benjamini-corrected p values < 0.05) for several GO categories and pathways in the *Fxn* knockdown co-expression modules which included several previously associated functional categories related with FRDA (**Supplementary Table 4**). Three modules (yellow, lightgreen and turquoise) that were down-regulated in two or in all three tissues due to *Fxn* knockdown included, nucleotide, nucleoside and ATP binding, myofibril assembly, muscle tissue development, RNA processing, and several mitochondrial related categories: oxidative phosphorylation, respiratory chain, NADH dehydrogenase activity, and electron transport chain. We also observed that the genes present in turquoise module were enriched for several KEGG pathways, namely, PPAR signaling (mmu03320; genes=14), insulin signaling (mmu04910; n=19), fatty acid metabolism (mmu00071; n=10), cardiac muscle contraction (mmu04260; n=20), dilated cardiomyopathy (mmu05414; n=13), and hypertrophic cardiomyopathy (mmu05410; n=14), which have been previously associated with FRDA, reflecting the multi-systemic nature of FRDA. Similarly, three upregulated cross tissue modules (blue, purple, and black) include, nucleotide binding, vesicle-mediated transport, immune response (innate and adaptive), defense response, inflammatory response, induction of apoptosis, positive regulation of cell death, cell adhesion, and skeletal system development (**Supplementary Table 4**). These results demonstrate that unsupervised analyses can identify groups of genes not only with shared biological functions, but also relevant to the clinical phenotypes observed in FRDA.

Three tissue specific modules that were down-regulated in heart and up-regulated in cerebellum (red, greenyellow and magenta) showed enrichment for, transcription regulator activity, neurological system process, synaptic vesicle and nucleotide, nucleoside and ATP binding. Two other modules that were up-regulated in heart and down-regulated in cerebellum (cyan and pink) were enriched for cell cycle, cell division, mitosis and DNA replication (**Supplementary Table 4**). In summary, we observed several metabolic functional categories that were differentially expressed (up and down) due to *Fxn* knockdown. The modules consisting of mitochondrial and cardiac specific categories along with PPAR signaling, insulin signaling, fatty acid metabolism pathways were down regulated in all tissues. Likewise, the modules enriched for immune, apoptosis and cell death related categories we up-regulated in all tissues due to *Fxn* knockdown. Synaptic and transcription regulator activity functional categories were only up-regulated in cerebellum, whereas cell cycle, cell division, mitosis and DNA replication related functional categories were down-regulated in cerebellum and up-regulated in the heart. These general functional categories related to *Fxn* knockdown have been previously associated with altered function in FRDA patients(44, 54), suggesting that genes within these modules would make interesting candidate genes for follow up studies, because many of the genes have not been previously associated with FRDA pathology and the disease mechanism.

### Gene expression candidate biomarkers associated with frataxin knockdown

To identify candidate molecular targets and to better understand the molecular mechanism associated with *Fxn* knockdown, we first manually combined all the GO ontology terms (see above and **Supplementary Table 4**) that were enriched in the 11 modules into 26 broad functional categories based on GO slim hierarchy (**Supplementary Table 5**) and screened for co-expressed genes within each functional category in all three tissues (r > 0.5 and P-value < 0.05) over the time course. This allowed us to identify critical functional sub-categories that are up or down regulated due to frataxin knockdown and subsequently permitted us to detect differentially expressed candidate genes that are co-expressed within each functional category (**Fig. 6d; Supplementary Fig. 9**). For example, we show that immune, cell cycle and apoptosis related functional groups are up-regulated, whereas cardiac, macromolecule and mitochondrial related functional groups were down-regulated (Fig. 6d). In the immune category, we observed most prominent changes in complement activation pathway genes, namely, C3, C4B, C1QB, C1QC and Serping1. Interesting, we also observed that many of these genes were also up-regulated in peripheral blood mononuclear cells obtained from FRDA patients (**Supplementary Fig. 10**), suggesting the potential for complement activation to act as a biomarker for FRDA as previously suggested for other degenerative diseases(55). Similarly, we found several genes, for example, CACNA2D1, ABCC9 and HRC involved in normal cardiac function, to be down-regulated in heart tissue upon frataxin knockdown (Fig. 6d). CACNA2D1 is associated with Brugada syndrome, also known as sudden unexpected nocturnal death syndrome, a heart condition that causes ventricular arrhythmia(56). Mutations in ABCC9 gene can cause dilated cardiomyopathy(57) and a genetic variant in the HRC gene has been linked to ventricular arrhythmia and sudden death in dilated cardiomyopathy(58). These observations suggest that lower levels of frataxin causes dysregulation of multiple genes related to arrhythmia or cardiac failure, a primary cause of death in FRDA patients.

There has been accumulating evidence suggesting that apoptosis may be an important mode of cell death during cardiac failure(59). In agreement with this, we observed genes related to apoptosis were up-regulated after *Fxn* knockdown in FRDAkd mice heart (Fig. 6d), which has been previously associated with FRDA pathogenesis and reported in other *Fxn* deficiency models(24, 60, 61). In order to validate our network findings, we tested CASP8 protein levels(62), observing an increase in cleaved Caspase 8 protein levels in Tg + heart tissue compared with control mice (Fig. 6e). Next, we employed the TUNEL assay to detect apoptotic cells that undergo extensive DNA degradation during the late stages of apoptosis(63). However, we did not observe an increase in cell death in all tissues by TUNEL staining (Fig. 6f).

### Literature data extraction for candidate genes associated with frataxin knockdown

We next examined the phenotype-gene associations extracted by co-occurrence-based text-mining in an attempt to link FRDA disease phenotypes with genes. For this, we screened the literature for potential co-occurrence link association between the observed FRDAkd mice phenotypes and the genes that are differentially expressed after *Fxn* knockdown (Online Methods). Identifying potential biomarker candidates that are previously validated for certain phenotypes can provide insight into disease progression, pathogenesis and extremely valuable for assessing therapeutic options(64). We screened with the genes that are differentially expressed (FDR < 5%) and present in the co-expression modules associated with behavioral and pathological key-terms (Eg: ataxia; **Supplementary Table 6**) in the published literature. Interestingly, this analysis identified numerous genes in which mutations are known to cause Mendelian forms of ataxia namely, ANO10(65, 66), CABC1(67, 68) and POLG(69, 70) that were significantly down-regulated in the heart tissue and mildly down-regulated in the cerebellum after *Fxn* knockdown in Tg + animals (**Fig. 6g and Supplementary Fig. 11**). We also found three other genes that are differentially expressed due to *Fxn* knockdown, namely, CSTB(71), NHLRC1(72) and PMM2(73) all of which are associated with other disorders when mutated causes ataxia (Fig. 6g). These observations suggest that *Fxn* along with multiple other downstream candidates causes behavioral deficits in FRDAkd mice. Similarly, for cardiac phenotypes, we identified multiple genes related to cardiac fibrosis that were up-regulated in FRDAkd mice heart (Fig. 6g), including LGALS3(74), ICAM1(75) and TIMP1(76) were up-regulated (Fig. 6g). Genes related to iron regulation, included HFE(77), SLC40A1(77), HMOX1(78), TFRC(77) and GDF15(79) all of which are directly involved in hemochromatosis and iron overload (Fig. 6g). We also found previously associated genes related with muscle strength (E.g.: SRL, DCN), myelination (E.g.: PMP22, LGALS3, TLR2 and SIRT2) and several genes related to neuronal degeneration (E.g.: GRN and APP) to be dysregulated in FRDAkd mice (**Fig. 6g and Supplementary Fig. 11**), connecting this degenerative disorder with the molecular signaling pathways known to be causally involved in other such disorders.

Perhaps the most prominent such known pathway associated with neurodegeneration, was autophagy, since several autophagy-related genes (LAMP2, ATF4, TLR2, OPTN, MAPK14, SIRT2, ICAM1, LGALS3, DCN, RCAN1, GRN) were present in these disease phenotype-associated sub networks (Fig. 6g). Disruption of autophagy is also reported as altered in other *Fxn* deficiency models(24, 60, 61) and is associated with various stress conditions including mitochondrial dysfunction(80, 81). Autophagy is responsible for the recycling of long-lived and damaged organelles by lysosomal degradation(82). To validate our network findings, we utilized LC3-II as a marker for autophagy, showing autophagy activation in Tg + mice heart, but not in spinal cord tissue (Fig. 6e, h). This suggests activation of apoptosis (Fig. 6d, e) and autophagy (Fig. 6e, h) may therefore potentiate the cardiac dysfunction of Tg + mice. Next, by examining the expression levels of all these sub network genes (Fig. 6g) in the datasets of other FRDA related mouse models and patient peripheral blood mononuclear cells, we show that they are also differentially expressed in the same direction (Fig. 6i), suggesting these sub network genes can be ideal candidates for molecular biomarkers in FRDA. In summary, consistent with our behavioral, physiological and pathological findings, we show multiple candidate genes related to key degeneration related phenotypes to be altered in FRDAkd mice.

### Rescue of behavioral, pathological and molecular changes due to frataxin restoration

Our data show that mice carrying the dox inducible shRNA for *Fxn* develop normally until they are challenged with dox, which subsequently results in substantial FXN reduction and causes multiple behavioral and pathological features observed in patients with FRDA, including cardiac and nervous system dysfunction (Fig. 2-5). *Fxn* knockdown mice displayed remarkable parallels with many of the behavioral, cardiac, nervous system impairments along with physiological, pathological and molecular deficits observed in patients. At a general level FRDAkd mice displayed weight loss, reduced locomotor activity, reduced strength, ataxia and early mortality (Fig. 2). In the nervous system, FRDAkd mice showed abnormal mitochondria and vacuolization in DRGs, retinal neuronal degeneration, and reduced axonal size and myelin sheath thickness in the spinal cord(8, 9, 24, 25) (Fig. 5). With regards to cardiac dysfunction, FRDAkd mice exhibited conduction defects, cardiomyopathy, evidence of iron overload, fibrosis, and biochemical abnormalities that are commonly observed in patients(8, 9, 25) (Fig. 3,4). These features correspond to a phenotype of substantial multisystem, clinical disability consistent with moderate to severe disease after 3 months of frataxin deficiency and that lead to a 50% mortality rate at approximately 5 months in these mice.

Given that optimal therapy in patients with FRDA could be considered replenishment of FXN itself (e.g. via gene therapy(83) or improvement of *Fxn* transcription via small molecules(84)), we next asked how much of the behavioral, physiological, pathological and molecular phenotype(s) observed at this relatively severe stage of illness could be reversed following such “optimal” therapy. In this case, optimal therapy is return of normal FXN levels under endogenous regulation through relief of exogenous inhibition. Answering this question of reversibility is crucial for any clinical trial, whatever the mechanism of action of the therapy, since we currently have limited information as to what represents potentially reversible neurologic or cardiac phenotypes.

To address this, we compared two groups of mice, both transgenic and treated with dox for 12 weeks, at which time both groups show equivalent levels of substantial clinical features (**Supplementary Fig. 12**). In one, Tg + the dox is continued for another 12 or more weeks, and in the other Tg ± the dox is removed and the animals are followed. Restoration of FXN expression in *Fxn* knockdown mice (Tg ±) that had reached a level of substantial clinical dysfunction led to significant improvement in lifespan (no death until 20 weeks post dox removal) when compared to Tg + group which resulted in 90% mortality rate by 25 week post dox treatment (Fig. 2b). Rescue animals (Tg ±) also displayed rapidly improvement in: gait ataxia, body weight, muscle strength, locomotor activity, and balance on rotarod test over the ensuing 12 week period, to a point where the treated animals were not significantly different from controls on many tests (Fig. 2). Remarkably, we observed all of the six FRDA associated clinical phenotypes tested showed significant improvement, suggesting that FRDA-like neurological defects due to absence of the mouse *Fxn* gene can be rectified by delayed restoration of *Fxn* (Fig. 2).

We next sought to determine whether the observed reversible behavioral changes in FRDAkd mice are also accompanied by recovery of the physiological phenotype in FRDAkd mice heart, since, changes in physiology offer attractive therapeutic targets for symptomatic and preventive treatment of ataxia. Our results in Tg ± mice which received the dox for 12 weeks followed by 12 weeks dox removal displayed reversal of long QT interval phenotype, when compared to Tg + mice at both 12 and 24 weeks post dox treatment (Fig. 3a,b,e). We observed the ventricular and posterior wall thickening only at 24 weeks post dox treatment in Tg + animals (Fig. 3d,f,g), suggesting that long QT interval phenotype is a prominent early manifestation of disease that occurs before left ventricular wall thickness and show that correcting this aberrant physiology through activation of *Fxn* gene expression is a potential route to therapy.

One question that intrigued us because of its striking behavioral and physiological functional recovery is to what extent these changes represented pathological findings related to cell dysfunction (potentially reversible) versus cell death (irreversible) recovery. Pathological and biochemical analyses in Tg ± mice heart at post 8 weeks dox withdrawal revealed improved cardiac function displaying reduced iron and ferritin accumulation, myocardial fibrosis, well ordered sarcomers, normal aconitase activity and reduced mitochondrial degeneration (Fig. 4). Relevant to the pathogenesis of FRDA heart and the role of iron and mitochondrial defect, it has been found that cells with these defects are sensitized to cellular dysfunction(85, 86), and here we show this can be ameliorated by *Fxn* restoration.

In the nervous system of Tg ± mice post 8 weeks dox removal, we observed reduced vacuoles and fewer condensed, degenerating mitochondria in DRG neurons along with several abnormal mitochondria containing DRG neurons (Fig. 5a,b,d,e). However we only observed mild improvement in myelin sheath thickness and cross section axonal size in the spinal cord of Tg ± mice during this time period (Fig. 5f,g). Conversely, we observed a significant reduction in the number of vacuoles and disrupted photoreceptors in the retina of Tg ± mice, indicating that *Fxn* restoration rescued photoreceptor degeneration (Fig. 5h,i). These findings establish the principle of cellular dysfunction reversibility in FRDAkd mouse model due to *Fxn* restoration and, therefore, raise the possibility that some neurological and cardiac defects seen in this model and FRDA patients may not be permanent.

In line with remarkable recovery of several behavioral, physiological and pathological defects in FRDAkd mice, we also observed that the genome-wide molecular biomarker represented by gene expression analyses due to *Fxn* knockdown could be completely rescued after *Fxn* restoration (Fig. 6). By rescuing the FXN protein levels back to the near basal level, we were able to reverse the molecular changes completely. At post 8 weeks of dox removal after initial 12 weeks of dox treatment, we examined the number of differentially expressed genes at false discovery rate of 5% in all tissues and observed 100% recovery of gene expression levels in cerebellum and DRGs, and 99.95 % of reversed gene expression levels in the heart of FRDAkd mice (Fig. 6a,b). These results which included several pathways (Fig. 6a,c,d) that are significantly affected due to *Fxn* depletion and complete reversal (Fig. 6a,b) of these pathways due to *Fxn* restoration can serve as gene-expression signature for evaluation of various therapeutic paradigm.

In summary, all six behavioral deficits in FRDAkd mice were reversed (Fig. 2), in the cardiac system, the long QT interval phenotype along with various pathological mainfestations including mitochondrial defects were reversed, in the nervous system, we observed improvement in DRGs and photoreceptor neurons, and complete reversal of the molecular changes in all three tissues suggesting near basal *Fxn* levels are sufficient to elevate behavioral symptoms in the preclinical FRDAkd model. The rapidity of *Fxn* expression due to dox removal and its robust correction of various parameters, even when restored after the onset of motor dysfunction, makes this FRDAkd mouse model an appealing potential preclinical tool for testing various therapeutics for FRDA.

## DISCUSSION

Here we report development and extensive characterization of a novel, reversible mouse model of FRDA based on knock-down of frataxin by RNA interference(28), the FRDAkd mouse. Treated transgenic mice developed abnormal mitochondria, exhibit cardiac and nervous system impairments along with physiological, pathological and molecular deficits. These abnormalities likely contributed to the behavioral phenotype of FRDAkd mice, parallel to what is observed in FRDA patients(8, 9). Importantly, restoration of *Fxn* expression, even after the development of severe symptoms and pathological defects, resulted in a drastic amelioration of the clinical phenotype, both in the cardiac and nervous systems, including motor activity. Pathology was significantly improved in the heart, DRGs and in the retina, but only mild improvement was observed in spinal cord, suggesting *Fxn* restoration for the duration of 8 weeks is not sufficient for reversal of neuronal defects in the spinal cord after *Fxn* knockdown. Interestingly, the patterns of gene expression changes due to *Fxn* knockdown in three different tissues differed(15), indicating variable response to frataxin deficiency, which could partly explain cell and tissue selectivity in FRDA(87). The onset and progression of the disease correlates with the concentration of doxycycline, and the phenotype returns to baseline after its withdrawal. The onset of transgene expression to achieve *Fxn* knockdown and robust recovery of symptoms due to restoration of *Fxn* levels in a mouse model of FRDA, even when reversed after the onset of the disease, makes this model an appealing potential preclinical tool for developing FRDA therapeutics. This approach will also enable new insights into FRDA gene function and molecular disease mechanisms.

Several models of FRDA have been developed and each have advantages and disadvantages(25). This new FRDAkd model exhibits several unique features that provide advantages for the study of FRDA pathophysiology relative to other existing models(20-25). First, induction of frataxin knockdown permits circumventing potential confounding developmental effects(19) and has the flexibility to enhance the disease onset and progression very rapidly by increasing the doxycycline dose. Moreover, the temporal control of *Fxn* knockdown can provide further insights into the sequence of tissue vulnerability during the disease progression(87). Second, reversibility of *Fxn* knockdown provides a unique model to mimic the effect of an ideal therapeutic intervention. Along this line, it has so far not been established to what extent FXN restoration after the onset of clinical motor symptoms is successful to prevent occurrence and/or progression of FRDA. Therefore, this model will be of central importance to gain better insights into disease pathogenesis and to test therapeutic agents. Third, the doxycycline-dependent reduction of *Fxn* expression can be tailored to carefully determine the critical threshold of *Fxn* levels necessary to induce selective cellular dysfunction in the nervous system (DRGs and spinal cord) and to understand the occurrence of tissue specific dysfunction in FRDA. Thereby, these experiments will help to understand the tissue specificity and generate clinically relevant tissue targeted therapeutics for FRDA. Finally, the temporal control of single copy and reversible regulation of shRNA expression against *Fxn* produces reproducible transgene expression from the well-characterized rosa26 locus to generate the first model to exhibit and reverse several symptoms parallel to FRDA patients. Here, we focused on the consequences of frataxin removal in an otherwise healthy adult animal. However, we should emphasize that this FRDAkd model uniquely facilitates future studies exploring prenatal or early post natal knockdown, and a wide variety of time course studies to understand disease pathophysiology and to identify potential imaging, physiological, or behavioral correlates of likely reversible or non-reversible disabilities in patients.

We show that FRDAkd animals are defective in several behaviors and exhibit weight loss, reduced locomotor activity, reduced strength, ataxia and early mortality. All of these defects were significantly improved following *Fxn* restoration, approaching or reaching wild-type levels. Most importantly ataxia and survival are well-established and important clinical endpoints in FRDA(31), readouts after *Fxn* restoration clearly improve these parameters and appear to be directly related to the functional status of the FRDAkd mice. We conclude that *Fxn* deficient mice exhibit considerable neurological plasticity even in a nervous system that is fully adult. Hence, utilizing these reversible intermediate behavioral phenotypes as biomarkers will help us determine the disease progression and test various FRDA treatment options in this model.

Hypertrophic cardiomyopathy is a common clinical feature in FRDA and approximately 60% of patients with typical childhood onset FRDA die from cardiac failure(31). It is generally believed that cardiac failure is caused by the loss of cardiomyocytes through activation of apoptosis(88). We observed activation of early apoptosis pathways in heart tissue and severe cardiomyopathy characterized by ventricular wall thickness(89). However, we did not observe TUNEL positive cells in either heart or nervous system. This may reflect that the model is in a early phase of cell death initiation, or rather that apoptotic cells are readily phagocytosed by neighbouring cells and are consequently difficult to detect(90). We also observed enhanced activation of autophagy in the heart tissue of FRDAkd mice, where autophagic cardiomyocytes are observed at a significantly higher frequency during cardiac failure(91). These results suggest that apoptosis and autophagy together might synergistically play a vital role in the development of cardiac defect in FRDA(92).

During *Fxn* knockdown, FRDAkd mice initially exhibited a long QT interval at 12 weeks during electrocardiographic analyses followed by the absence of P-waves and increased ventricular wall thickness at 24 weeks. Restoration of *Fxn* levels at 12 weeks reversed long QT interval phenotype. However, it will be interesting to examine if the ventricular wall thickness can be restored by a more prolonged rescue time period. Another prominent feature of *Fxn* deficiency mouse and FRDA patients is iron accumulation and deficiency in activity of the iron-sulfur cluster dependent enzyme, aconitase, in cardiac muscle(23, 40, 85, 86). Consistent with these observations, we observed increased iron accumulation and reduced aconitase activity in the cardiac tissue of FRDAkd mice and we demonstrate a marked reversal of both to a statistically significant extent, suggesting *Fxn* restoration is sufficient to overcome and clear the iron accumulation and reverse aconitase activity(93). Our gene expression data revealed several genes (HFE(77), SLC40A1(77), HMOX1(78), TFRC(77) and GDF15(79)) directly involved in hemochromatosis and iron overload to be upregulated in our FRDAkd mice, all of which were rescued to normal levels by frataxin restoration. Similarly, several downregulated genes involved in normal cardiac function (CACNA2D1, ABCC9 and HRC) were rescued by *Fxn* restoration. Together, these data indicate that *Fxn* restoration in symptomatic FRDAkd mice reverses the early development of cardiomyopathy at the molecular, cellular and physiological levels.

Cellular dysfunction due to FXN deficiency is presumed to be the result of a mitochondrial defect, since FXN localizes to mitochondria(93-95) and deficiencies of mitochondrial enzymes and function have been observed in tissues of FRDA patients(40, 96). Here we show that FRDAkd mice displayed accumulation of damaged mitochondria, and reduced aconitase as a direct consequence of frataxin deficiency in heart, consistent with previous findings in conditional FRDA mouse models(23, 24), suggesting frataxin deficiency inhibits mitochondrial function leading to cellular atrophy(97). Restoration of FXN resulted in improvement in pathological mitochondrial structure indicating that FXN restoration prevents mitochondrial defects and may thereby enhance cell survival(93). Our observation of mitochondrial dysfunction recovery as early as eight weeks after FXN restoration is consistent with the highly dynamic nature of mitochondria(98).

In the nervous system, we observed higher level of condensed mitochondria in DRGs through ultrastructural analyses. Condensed mitochondria are known to have lower respiratory control and ATP production(99). On most occasions these condensed mitochondria in DRG neurons of FRDAkd mice were associated with lipid-like bodies, consistent with the known close association of lipid bodies with mitochondria observed in a variety of cell types(100). Of particular interest, it has been shown that the junctions between these two organelles expand with an increased need for energy, suggesting the direct flow of fatty acids from lipid bodies to the mitochondrial matrix for β-oxidation to meet the cell’s energy needs(101). We observed reduced condensed mitochondria and their association with lipid-like bodies in rescue animals, suggesting that a substantial fraction of dysfunctioning frataxin-deficient mitochondria containing neurons are still viable after the onset of disease and that their dysfunction can be reversed. In the spinal cord, we observed reduction of axonal size and myelin sheath thickness in FRDAkd animals, however after eight weeks of rescue period by FXN restoration, we observed limited improvement, suggesting more time may be necessary for improved nervous system recovery. Conversely, disruption of photoreceptor neurons and degenerating RPE cells in the retina displayed robust recovery, indicating an overall morphological improvement in retinal neurons upon FXN restoration to normal levels. These findings extend previous studies showing that many defects in FRDA cells in vitro and cardiac function in vivo may be reversible following reintroduction of FXN(102) by pharmacological interventions targeting aberrant signaling processes(103) or by reintroduction of FXN by gene therapy(83).

Improved understanding of the mechanism by which FXN deficiency leads to the various phenotypes observed in FRDA is a major goal of current research in this disorder. In this regard, gene expression analyses identified several pathways altered, including, PPAR signaling pathway, insulin signaling pathway, fatty acid metabolism, cell cycle, protein modification, lipid metabolism, and carbohydrate biosynthesis, all of which have been previously associated with altered function in FRDA patients(15, 54, 104). We also observed several immune components, namely, complement and chemokine cascade genes being upregulated, may be implicated as a protective mechanism after *Fxn* knockdown, and in a causal role through chronic activation of the inflammatory response(105, 106). It will be interesting to validate these genes and pathways as potent candidate biomarkers for FRDA at several stages during the disease progression.

Determining definitively whether the global expression changes observed in this study are primary or secondary to *Fxn* knockdown will require further investigation by examining dense time points immediately after *Fxn* knockdown and rescue, but our data clearly support the utility of induced models of severe disease, whereby the consequences of gene depletion can be more cleanly controlled to examine the molecular changes(28). Together with post-induction rescue, this study highlights potential biomarkers and pathways of FRDA progression. For FRDA clinical trials, it will be important to assess whether these expression changes are translatable to the human disease and extend to other tissues that are more easily accessible than CNS or cardiac tissue. Together, these observations suggest that this work also provides a functional genomics foundation for understanding FRDA disease mechanism, progression and recovery.

In conclusion, our study provides multiple lines of evidence that *Fxn* knockdown in adult mice leads to several symptoms parallel to FRDA patients. By restoring FXN levels we show reversal of several symptoms even after significant motor defect, demonstrating the utility of this attractive model for testing potential therapeutics, such as gene therapy(83), protein replacement therapy(102), enhancement of mitochondrial function(107) and small molecules(108, 109). In fact, our findings suggest that such approaches may not only enhance FXN expression and rescue downstream molecular changes, but may too alleviate pathological and behavioral deficits associated with FRDA, depending on the stage of the disorder.

## METHODS

### In vitro frataxin knockdown

Six different shRNA sequence against the frataxin mRNA (**Supplementary Fig. 1**) were cloned into the exchange vector (proprietary material obtained from TaconicArtemis GmbH) as described in detail previously(28). N2A cells (ATCC) were transduced with exchange vectors containing the shRNA against frataxin using Fugene 6 (Promega, Madison, WI). After 24 h, cells were replated in media containing neomycin and were validated for efficient frataxin knockdown by RT-PCR and Western blotting. Mouse LW-4 (129SvEv) embryonic stem cells were cultured and the rosa26 targeting vector (from TaconicArtemis GmbH) were electroporated according to protocol described before(12). The exchange vector containing validated and selected shRNA sequence (GGATGGCGTGCTCACCATTAA) against the Fxn mRNA were electroporated to obtain positive ES cells containing shRNA expression cassette integrated into the ROSA26 locus. Correctly targeted embryonic stem cell clones were utilized to generate frataxin knockdown mice (see below).

### Frataxin knockdown transgenic mice generation

Transgenic mice were generated at UCLA Transgenic Core facility using proprietary materials obtained from TaconicArtemis GmbH (Köln, Germany). In brief, mouse LW-4 (129SvEv) embryonic stem cells with recombinase-mediated cassette exchange (RMCE) acceptor site on chromosome 6 were used for targeting insertion of distinct Tet-On frataxin shRNA expression cassettes into the ROSA26 locus(28) as depicted in Fig. 1. Correctly targeted embryonic stem cell clones were identified by Southern DNA blot analysis and tested for frataxin mRNA knockdown at the embryonic stem cell stage (not shown). One embryonic stem cell clone that gave acceptable mRNA knockdown were microinjected into C57BL/6J blastocysts from which chimeric mice were derived. Frataxin knockdown mice were backcrossed six generations into C57BL/6 mice.

### Genotyping

Mouse tail biopsies were collected and DNA was extracted in boiling buffer (25 mM sodium hydroxide, 0.2 mM EDTA) at 98°C for 60 min. Extracted DNA was neutralized in Tris/HCl buffer (pH5.5) and PCR was performed under the following conditions with BioMix Red (Bioline). Four primers were used in the reaction: 5’-CCATGGAATTCGAACGCTGACGTC-3’, 5’-TATGGGCTATGAACTAATGACCC-3’, to amplify shRNA; 5’-GAGACTCTGGCTACTCATCC-3’, 5’- CCTTCAGCAAGAGCTGGGGAC-3’, as genomic control. The cycling conditions for PCR amplification were: 95°C for 5 min; 95°C for 30 s, 60°C for 30 s, 72°C for 1 min, (35 cycles); 72°C for 10 min. PCR products were analyzed by gel electrophoresis using 1.5% agarose and visualized by Biospectrum imaging system (UVP).

### Animal and study design

Experiments were approved by the Animal Research Committee (ARC) of University of California, Los Angeles and were in accordance with ARC regulation. Extensive neurological and neuropsychological tests (body weight, poorly groomed fur, bald patches in the coat, absence of whiskers, wild-running, excessive grooming, freezing, hunched body posture when walking, response to object [cotton-tip swab test], visual cliff behavior analysis) were performed to ensure all animals included in the studies were healthy. Age and sex were matched between wild type (Wt) and transgenic (Tg) groups to eliminate study bias. The average age of the animals at the start of experiments was 3-4 months. Three different study cohorts were implemented: behavior, pathology and gene expression. Animals were randomly assigned to different experimental groups with in these three different study cohorts before the start of experiments. Due to high mortality rate of FRDAkd animals, we doubled the Tg + animal number (a sample size of 30 animals per Tg + treatment group for behavioral analyses (60 total)) in order to have sufficient power for statistical analyses. All the group size for our experiments were determined by statistical power analysis. The values utilized were: the power = 0.9, alpha = 0.05, Coefficient of determination = 0.5, effect size = 0.70. Effect size and power calculations were based on our pilot experiments.

### In vivo frataxin knockdown

Animals were divided into the following groups: Wt treated with doxycycline (Dox) (Wt +), Wt without Dox (Wt -), Tg treated with Dox (Tg +), Tg without Dox (Tg -), Tg Dox rescue (Tg ±). First, we examined the Tg + mice for frataxin knockdown utilizing higher dose of dox (4 and 6mg/ml), we observed mortality as early as two weeks and a 100% mortality rate by 5 to 6 weeks (not shown). To avoid early mortality and to have slow and steady state of disease progression we followed 2 mg/ml in drinking water coupled with intraperitoneal injection of dox (5 or 10 mg/kg) twice per week. Doxycycline (2mg/mL) was added to the drinking water of all treatment animals which was changed weekly. In addition, animals were injected intraperitoneally (IP) with Dox (5 mg/kg body weight) twice a week for 10 weeks followed by 10 mg Dox/kg body weight twice a week for 2 weeks. Animals in Tg Dox rescue (Tg ±) group were given untreated water and not injected with Dox after week 12. All animals were weighed weekly.

### Behavior cohort

*Animal information.* A total of 108 animals (Wt n=32, Tg n=76) were included in this study with equal numbers of male and female animals. For all tests, investigators were blinded to genotype and treatment. For all behavioral tests, the variance between all the groups for that specific behavioral test were observed to be initially not statistically significant.

*Accelerating Rotarod.* To measure motor function, rotarod analysis was performed weekly at the start of Dox treatment using an accelerating rotorod (ROTOMEX-5, Columbus Instruments, Columbus, OH). Mice were assessed for 36 weeks. Briefly, after habituation, a mouse was placed on the rotarod rotating at 5 rpm for one min and then the rotorod was accelerated at 0.09 rpm/sec^2^. The latency to fall from a rotating rod after acceleration was recorded. Each mouse was subjected to 3 test trials within the same day with a 15 min inter-trial interval. The average latency normalized by the mouse body weight in the test week was used for data analysis.

*Grip strength test*. The grip strength was measured at week 0, 12 and 24 weeks using a digital force gauge (Chatillon Force Measurement Systems, AMETEK TCI Division, Largo, FL). Briefly, the mouse was allowed to grasp the steel wired grid attached to the force gauge with only forepaws. The mouse was then pulled back from the gauge. The force applied to the grip immediately before release of the grip is recorded as the “peak tension” and is a measurement of forepaw strength. The same measurement was repeated with the mouse grasping the grid with all paws for whole grip strength. Each mouse was subjected to three forepaw and whole strength measurements. The hindpaw grip strength was calculated as the average whole grip strength minus average forepaw strength. The value of hindpaw grip strength normalized by the mouse body weight in the test week was used for data analysis.

*Gait analyses*. Gait analysis was performed at week 0, 12 and 24 weeks by allowing the animals to walk through a 50-cm-long, 10-cm-wide runway that was lined with blank index cards. After a period of habituation (walking through the runway three times), hind and fore paws were coated with nontoxic red and purple paint respectively and the mouse was allowed to walk through the runway again. Footprints were captured on the index cards. The index cards were scanned and the images were measured in image J for calculating the stride length and other parameters.

*Open field analyses*. Open field testing was performed at week 0, 12 and 24 weeks. Mice were placed in a clear plexiglass arena (27.5 cm by 27.5 cm) with a video monitoring system that recorded any movements of the mouse within the chamber for 20 min. Locomotor activity of the mice was analyzed by TopScan (CleverSys) software.

### Pathology cohort

*Animal information.* A total of 57 animals (Wt n=21 (10 females, 11 males), Tg n=36 (20 females, 16 males)) were euthanized in the study. Animals were deeply anesthetized with sodium pentobarbital (40 mg/kg body weight) and perfused intracardially with 20 mL PBS (5 mL/min) followed by 20 mL freshly made 4% paraformaldehyde (5 mL/min). Tissue from liver, lung, spleen, pancreas, kidney, heart, eye (retina), brain, muscle, spinal cord, dorsal root ganglion (DRG) and sciatic nerve was dissected and collected immediately after perfusion. Collected tissue was immersed in 4% paraformaldehyde overnight at 4°C and then transferred and kept in 40% sucrose at 4°C until embedded with O.C.T. Tissue cryosections (5 µM) were collected with a cryostat for staining.

*Staining.* H & E staining was performed in Translational Pathology Core Laboratory at UCLA. The method is described briefly below. After equilibration, the tissue section was stained with Harris Modified Hematoxylin for 10 min, then in Eosin Y for 10 min. Gomori’s iron staining and Masson’s trichrome staining were performed in Histopathology lab in UCLA, the procedure is described briefly as below. Gomori’s iron staining: the tissue section was immersed for 20 min in equal parts of 20% hydrochloric acid and 10% potassium ferrocyanide, washed thoroughly in distilled water. After counterstaining with Nuclear Fast Red (0.1% Kernechtrot in 5% aluminum sulfate), the slides were dehydrated through gradual ethyl alcohol solutions and mounted for imagining. Masson’s trichrome staining: the tissue section was fixed in Bouin’s fixative (75 mL picric acid saturated aqueous solution, 25 mL of 40% formaldehyde, 5 mL glacial acetic acid) for 60 min at 60°C, stained with Weigert iron hematoxylin (0.5% hematoxylin, 47.5% alcohol, 0.58% ferric chloride, 0.5% hydrochloric acid) for 10 min, Biebrich scarlet-acid fuchsin (0.9% Biebrich scarlet, 0.1% acid fuchsin, 1% glacial acetic acid) for 10 min, immersed in phosphomolybdic-phosphotungstic solution (2.5% phosphomolybdic acid, 2.5% phosphotungstic acid) for 7 min, stained with aniline blue solution (2.5% aniline blue, 2% acetic acid) for 7 min, dehydrate and mounted for imagining. Immunofluorescence staining was performed as followed: cryosections were equilibrated with TBS (20 mM Tris pH7.5, 150 mM NaCl) and permeabilized in TBST (TBS with 0.2% TritonX-100). Tissue sections were incubated with primary antibody in TBST with 4% normal goat serum (Jackson Immuno Research) (4°C, overnight), followed by incubation with secondary antibody in TBST with 4% normal goat serum (room temperature, 2 hrs) the next day. The tissue section was mounted with ProLong Gold with DAPI (ThermoFisher Scientific) and stored in the dark. The following antibodies were used: anti-ferroportin-1 (LifeSpan BioSciences, LS-B1836, rabbit, 1:200), anti-ferritin (Abcam, ab69090, rabbit, 1:500), anti-LC3 (Abgent, AM1800a, mouse, 1:200), anti-NeuN (EMD Millipore, ABN91, chicken, 1:500), anti-NeuN (EMD Millipore, ABN78, rabbit, 1:500). Secondary antibodies conjugated with Alexa Fluor (ThermoFisher Scientific, 1:500) were used as indicated: Alexa Fluor 488 (goat anti-Rabbit IgG, A-11008), Alexa Fluor 594 (goat anti-chicken IgG, A-11039), and Alexa Fluor 488 (goat anti-mouse IgG, A-11029). DNA strand breaks were determined by TUNEL assay (Roche). Briefly, sections were equilibrated with PBS, permeabilized in 0.1% Triton X-100 in 0.1% sodium citrate (on ice, 2 min) and incubated with TUNEL reaction mixture in a humidified atmosphere (37°C, 60 min). Sections were mounted with ProLong Gold with DAPI and stored in the dark.

*Imaging and quantification.* Slides stained for hematoxylin-eosin, iron, and trichrome were scanned (20 x magnification) by Aperio ScanScope with Aperio ImageScope Software (Leica). Immunofluorescence staining and TUNEL staining were imaged with LSM780 confocal microscope system (Zeiss). Images were quantified by CellProfiler. For Purkinje cell quantification, the H&E images were scanned, visualized and exported by ImageScope (Leica Biosystems) system. Exported images were utilized to count the Purkinje cells manually using NIH ImageJ software. Counters were blinded to genotype and treatment.

*Electron microscopy analyses.* A total of 29 animals (Wt n=9 (4 females, 5 males), Tg n=20 (9 females, 11 males)) were used in the study. Mice were perfused transcardially with 2.5% glutaraldehyde, 2% paraformaldehyde in 0.1M phosphate buffer, 0.9% sodium chloride (PBS). Pieces of heart, lumbar spinal cord, dorsal root ganglia, muscle, and eye were dissected, postfixed for 2 hours at room temperature in the same fixative and stored at 4°C until processing. Tissues were washed with PBS, postfixed in 1% OsO4 in PB for 1 hour, dehydrated in a graded series of ethanol, treated with propylene oxide and infiltrated with Eponate 12 (Ted Pella) overnight. Tissues were embedded in fresh Eponate, and polymerized at 60°C for 48 hours. Approximately 60-70 nm thick sections were cut on a RMC Powertome ultramicrotome and picked up on formvar coated copper grids. The sections were stained with uranyl acetate and Reynolds lead citrate and examined on a JEOL 100CX electron microscope at 60kV. Images were collected on type 4489 EM film and the negatives scanned to create digital files. These high quality digital images were utilized to quantify the number of condensed mitochondria. Condensed mitochondria, vacuoles with condensed mitochondria, and vacuoles alone were manually counted with NIH ImageJ software. Animal genotype and treatment information was blinded to the person who conducted the evaluation.

### Gene expression cohort

*Sample collection.* A total of 80 animals (Wt n=24 (12 females and 12 males), Tg n=56 (31 females and 25 males)) were sacrificed and tissue was collected in this study. Animals were sacrificed at week 0, 3, 8, 12, 16, 20 and plus 4, 8 weeks post dox treatment (rescue). Mouse was sacrificed by cervical dislocation and tissue from liver, lung, spleen, pancreas, kidney, heart, eye (retina), brain, muscle, spinal cord, dorsal root ganglion (DRG) and sciatic nerve was dissected and rinsed in cold PBS quickly (3X) to remove blood. Tissue samples were transferred immediately into 2 mL RNase-free tubes and immersed into liquid nitrogen. The collected tissue was stored in -80°C freezer immediately.

*RNA extraction.* Heart, cerebellum and DRG neuron samples from week 0, 3, 12, 16, 20 and 4, 8 weeks post dox treatment (rescue) with four biological replicates were used for expression profiling. Samples were randomized prior to RNA extraction to eliminate extraction batch effect. Total RNA was extracted using the miRNeasy mini kit (Qiagen) according to manufacturer’s protocol and including an on-column DNase digest (RNase free DNAse set; Qiagen). RNA samples were immediately aliquoted and stored at -80°C. RNA concentration and integrity were later determined using a Nanodrop Spectrophotometer (ThermoFisher Scientific) and TapeStation 2200 (Agilent Technologies), respectively.

*Transcriptome profiling by microarray*. One hundred nanograms of RNA from heart and cerebellum tissue was amplified using the Illumina TotalPrep-96 RNA Amplification kit (ThermoFisher Scientific) and profiled by Illumina mouse Ref 8 v2.0 expression array chips. For DRG samples 16.5 ng of RNA was amplified using the Ovation^®^ PicoSL WTA System V2 kit (NuGEN). Only RNA with RIN greater than 7.0 was included for the study. A total of 64 RNA samples for each tissue (n= 192 arrays) were included and samples were randomized before RNA amplification to eliminate microarray chip batch effect. Raw data was log transformed and checked for outliers. Inter-array Pearson correlation and clustering based on variance were used as quality-control measures. Quantile normalization was used and contrast analysis of differential expression was performed by using the LIMMA package(32). Briefly, a linear model was fitted across the dataset, contrasts of interest were extracted, and differentially expressed genes for each contrast were selected using an empirical Bayes test statistic(32).

*Construction of co-expression networks.* A weighted gene co-expression network was constructed for each tissue dataset to identify groups of genes (modules) associated with temporal pattern of expression changes due to frataxin knockdown and rescue following a previously described algorithm(49, 110). Briefly, we first computed the Pearson correlation between each pair of selected genes yielding a similarity (correlation) matrix. Next, the adjacency matrix was calculated by raising the absolute values of the correlation matrix to a power (β) as described previously(49). The parameter β was chosen by using the scale-free topology criterion(49), such that the resulting network connectivity distribution best approximated scale-free topology. The adjacency matrix was then used to define a measure of node dissimilarity, based on the topological overlap matrix, a biologically meaningful measure of node similarity(49). Next, the probe sets were hierarchically clustered using the distance measure and modules were determined by choosing a height cutoff for the resulting dendrogram by using a dynamic tree-cutting algorithm(49).

*Consensus module analyses.* Consensus modules are defined as sets of highly connected nodes that can be found in multiple networks generated from different datasets (tissues). Consensus modules were identified using a suitable consensus dissimilarity that were used as input to a clustering procedure, analogous to the procedure for identifying modules in individual sets as described elsewhere(111). Utilizing consensus network analysis, we identified modules shared across different tissue data sets after frataxin knockdown and calculated the first principal component of gene expression in each module (module eigengene). Next, we correlated the module eigengenes with time after frataxin knockdown to select modules for functional validation.

*Gene Ontology, pathway and PubMed analyses.* Gene ontology and pathway enrichment analysis was performed using the DAVID platform (DAVID, https://david.ncifcrf.gov/(112)). A list of differentially regulated transcripts for a given modules were utilized for enrichment analyses. All included terms exhibited significant Benjamini corrected P-values for enrichment and generally contained greater than five members per category. We used PubMatrix(113) to examine each differentially expressed gene’s association with the observed phenotypes of FRDAkd mice in the published literature by testing association with the key-words: ataxia, cardiac fibrosis, early mortality, enlarged mitochondria, excess iron overload, motor deficits, muscular strength, myelin sheath, neuronal degeneration, sarcomeres, ventricular wall thickness, and weight loss in the PubMed database for every gene.

*Data Availability.* Datasets generated and analyzed in this study are available at Gene Expression Omnibus. Accession number: GSE98790.

### Quantitative real-time PCR

RT-PCR was utilized to measure the mRNA expression levels of frataxin in order to identify and validate potent shRNA sequence against frataxin gene. The procedure is briefly described below: 1.5 µg total RNA, together with 1.5 µL random primers (ThermoFisher Scientific, catalog# 48190-011), 1.5 µL 10 mM dNTP (ThermoFisher Scientific, catalog# 58875) and RNase-free water up to 19.5 µL, was incubated at 65 °C for 5 min, then on ice for 2 min; 6 µL first strand buffer, 1.5 µL 0.1 M DTT, 1.5 µL RNaseOUT (ThermoFisher Scientific, catalog# 100000840) and 1.5 µL SuperScript III (ThermoFisher Scientific, catalog# 56575) were added to the mixture and incubated for 5 min at 25 °C, followed by 60 min at 50 °C, and 15 min at 70 °C. The resulted cDNA was diluted 1:10 and 3 µL was mixed with 5 µL SensiFast SYBR No-ROX reagent (Bioline, catalog# BIO-98020), 0.6 µL primer set (forward and reverse, 10 mM) and 1.4 µL nuclease free water for real-time PCR. Each reaction was repeated three times on LightCycler 480 II (Roche, catalog# 05015243001). The thermocycling program is as following: 95 °C 5min; 95 °C 10 sec, 60 °C 10 sec, 72 °C 10 sec, repeat 45 cycles; 95 °C 5 sec, 65 °C 1 min, 97 °C 5 min.

### Western blotting

Prior to samples being analyzed by Western blotting, protein concentration was estimated by Bradford assay (Bio-Rad). Proteins were separated by SDS-PAGE and transferred onto PVDF membranes. The membranes were incubated with primary and secondary antibodies for protein detection. The target bands were detected with antibodies: anti-Frataxin (Santa Cruz Biotechnology, sc-25820, rabbit, 1:200), anti-Akt (Cell Signaling Technology, #4691, rabbit, 1:500), anti-β-Actin (Sigma, #A1978, mouse, 1:1000), anti-Caspase 8 (rabbit monoclonal, Cell Signaling Technology, #8592S, rabbit, 1:200), anti-LC3 (Abgent, # AM1800a, mouse, 1:200) and goat-anti-rabbit or goat-anti-mouse HRP-conjugated secondary antibodies (ThermoFisher Scientific, #31460, and #32230, respectively,1:5000). Image J was used for band quantification.

### Enzyme activity assay

Proteins from heart tissue for all genotypes at week 20 was extracted in Tris buffer (50 mM Tris-HCl pH 7.4). Around 80 mg of tissue was homogenized with 100 μL Tris buffer, and centrifuged at 800 g for 10 min at 4°C. The supernatant was transferred to a clean tube and stored at -80°C freezer immediately for later use. Protein concentration was estimated by Bradford assay. Total of 5 μg of protein was used to determine aconitase activity by aconitase assay kit (Cayman Chemical, catalog# 705502). The absorbance of the reaction mixture at 340 nm was measured every minute for 60 min at 37°C, by Synergy-2 (BioTek) plate reader. The linear phase was used to calculate the enzyme activity. 5 μg of protein was used to measure citrate synthase activity by citrate synthase assay kit (Sigma, catalog# CS0720). The absorbance of the reaction mixture at 412 nm was measured by Synergy-2 plate reader every 31 seconds for 30 min at room temperature. The linear phase was used to calculate the enzyme activity. Aconitase activity in each sample was normalized to the citrate synthase activity from the same sample for comparison.

### Echocardiography

Echocardiography was performed on all the mice with a Siemens Acuson Sequoia C256 instrument (Siemens Medical Solutions, Mountain View, California). The mice were sedated with isoflurane vaporized in oxygen (Summit Anesthesia Solutions, Bend Oregon). Left ventricular dimensions (EDD, end diastolic dimension; ESD, end systolic dimension; PWT, posterior wall thickness; VST, ventricular septal thickness) were obtained from 2D guided M-mode and was analyzed using AccessPoint software (Freeland System LLC, Santa Fe, New Mexico) during systole and diastole. Left ventricular mass was calculated as described previously(18). Heart rate, aortic ejection time, aortic velocity and mitral inflow E & A wave amplitudes were determined from Doppler flow images. Indices of contractility such as left ventricular fractional shortening (LVFS), velocity of circumferential fiber shortening (VCF) and ejection fraction (EF) was obtained from the images. During the procedure, heart rates were maintained at physiological levels by monitoring their electrocardiograms (ECG).

### Surface ECG

Electrocardiogram (ECG) were obtained in all the mice under Isoflurane anesthesia by inserting two Pt needle electrodes (Grass Technologies, West Warwick, RI) under the skin in the lead II configuration as described previously(27). The mice were studied between 10 and 30 minutes to elucidate any rhythm alteration. The ECG data were amplified (Grass Technologies) and then digitized with HEM V4.2 software (Notocord Systems, Croissy sur Seine, France). Measurements of HR, PR, RR, QRS, QT intervals were obtained for comparison and statistical analysis among all the mice groups.

**Accession codes.** Gene Expression Omnibus: Datasets generated and analyzed in this study are available at GEO accession: GSE98790.

## ACKNOWLEDGMENTS

We gratefully acknowledge support from the Friedreich’s Ataxia Research Alliance to D.H.G. and V.C. (including a New Investigator Award to V.C.), and the Dr. Miriam and Sheldon G. Adelson Medical Research Foundation to D.H.G.. We thank Marianne Cilluffo for assistance with electron microscopy at the BRI Electron Microscopy Core Facility at UCLA.

## AUTHOR CONTRIBUTIONS

V.C. and D.H.G. designed the experiments and wrote the manuscript. K.G., R.V., H.D., and V.C. performed the behavioral, pathological, biochemical and molecular experiments. V.C. conducted or coordinated all the experiments discussed in this manuscript with input from D.H.G.. V.S. performed network analyses and V.C. performed all subsequent bioinformatic analyses. M.J. and V.C. conducted and coordinated electrocardiogram and echocardiogram analyses. All authors discussed the results and provided comments and revisions on the manuscript.

